# Hepatocyte Cell Cycle Progression Depends on a Transcriptional Repressor Cascade Downstream of Wnt Signaling

**DOI:** 10.1101/2021.10.15.464616

**Authors:** Yinhua Jin, Teni Anbarchian, Peng Wu, Abby Sarkar, Matt Fish, Roel Nusse

## Abstract

Cell proliferation is tightly controlled by inhibitors that block cell cycle progression until growth signals relieve this inhibition. In several tissues including the liver, transcriptional repressors such as E2F7 and E2F8 function as inhibitors of mitosis and promote polyploidy, but how growth factors release these mitotic inhibitors to facilitate cell cycle progression is unknown. We describe here a newly identified mechanism of cell division control in which Wnt/βcatenin signaling in the postnatal liver maintains active hepatocyte proliferation through *Tbx3*, a Wnt target gene. TBX3 directly represses transcription of *E2f7* and *E2f8*, promoting a low ploidy state and cell cycle progression. This sequential transcriptional repressor cascade, initiated by Wnts, provides a new paradigm for exploring how a commonly active developmental signal impacts cell cycle completion.

## Introduction

Cells in adult tissues mostly remain quiescent unless they become activated by growth factors to enter the cell cycle and proliferate (Jones and Kazlauskas 2001). Quiescence is commonly imposed by inhibitors that prevent cells from entering S phase (Manning and Cantley 2007). These inhibitors are in turn repressed by growth factors that allow cell cycle entry by activating the G1/S transition (Coller 2007). In some tissues including the liver, cells can enter S phase to duplicate their genome but do not complete M phase thus becoming polyploid (Donne et al. 2020; Guidotti et al. 2003; Øvrebø and Edgar 2018). Two members of the E2F family of cell cycle regulators, E2F7 and E2F8, are responsible for this form of cell cycle arrest in the liver (Chen et al. 2012; Pandit et al. 2012). Acting as mutually redundant transcriptional repressors, E2F7 and E2F8 inhibit the expression of many genes, including *Aurka/b, Ccnb1* and *Plk1*, which encode proteins that act during mitotic progression (Chen et al. 2012; de Bruin et al. 2003; Pandit et al. 2012). Deleting *E2f7* and *E2f8* from these cells results in completion of M phase and the generation of diploid daughter cells (Chen et al. 2012; Pandit et al. 2012). Therefore, in the liver, DNA synthesis is uncoupled from cell proliferation due to the activity of E2F7 and E2F8. Whether growth factors can suppress the activity of E2F7 and E2F8 and thereby promote cell division is currently not known.

The postnatal liver expands rapidly in a Wnt/βcatenin signaling-dependent way and switches from initially containing diploid hepatocytes to becoming mostly polyploid (Apte et al. 2007; Monga et al. 2001). Here, we have studied the regulation of hepatocyte cell division *in vivo* as well as in primary cell culture to elucidate the molecular mechanisms of cell cycle progression. In the mouse and human liver, Wnt ligands are produced by endothelial cells of the central vein branches and the nearby sinusoids (McEnerney et al. 2017; Wang et al. 2015; Zhao et al. 2019), creating a zonated expression pattern of Wnt target genes and key metabolic enzymes in pericentral hepatocytes (Halpern et al. 2017). The transcriptional repressor *Tbx3*, a gene required for embryonic liver development (Khan et al. 2020; Ludtke et al. 2009; Suzuki et al. 2008), is also expressed in a zonated pattern during postnatal liver growth (fig. S1A) (Wang et al. 2015; Zhao et al. 2019).

## Results

### Tbx3 promotes hepatocyte proliferation

We verified that *Tbx3* is a target of Wnt signaling in the liver, by either over-activating or eliminating the Wnt pathway in the tissue (Figure. 1A, arrows point to TBX3^+^ *Apc* KO cells, arrowheads point to TBX3^-^ *Wls* KO cells), in agreement with data from other contexts (Aydoğdu et al. 2018; Eblaghie et al. 2004; Renard et al. 2007; Waghray et al. 2015). To examine the function of TBX3 in hepatocytes, we employed a recently developed method for long-term culture and genetic manipulation of primary hepatocytes, where cells multiply rapidly in a Wnt-dependent manner (Peng et al. 2018). We knocked down *Tbx3* in cultured hepatocytes using two different shRNA constructs (Supplementary Figure 1B), which significantly slowed down hepatocyte proliferation (Figure 1B) and led to increased numbers of polyploid cells (Figure 1C, Figure S1C and Table S1), suggesting that hepatocytes failed to complete M-phase. Conversely, overexpression of *Tbx3* in hepatocytes was sufficient to increase proliferation (Figure 1D and Figure S1, D and E), even in the absence of CHIR-activated Wnt signaling (Figure 1D) and increased the percentage of diploid cells (Figure 1E and Table S1). As expected, CHIR alone enhanced proliferation of hepatocytes albeit not to the same degree as *Tbx3* overexpression (Figure 1D).

**Figure 1.**
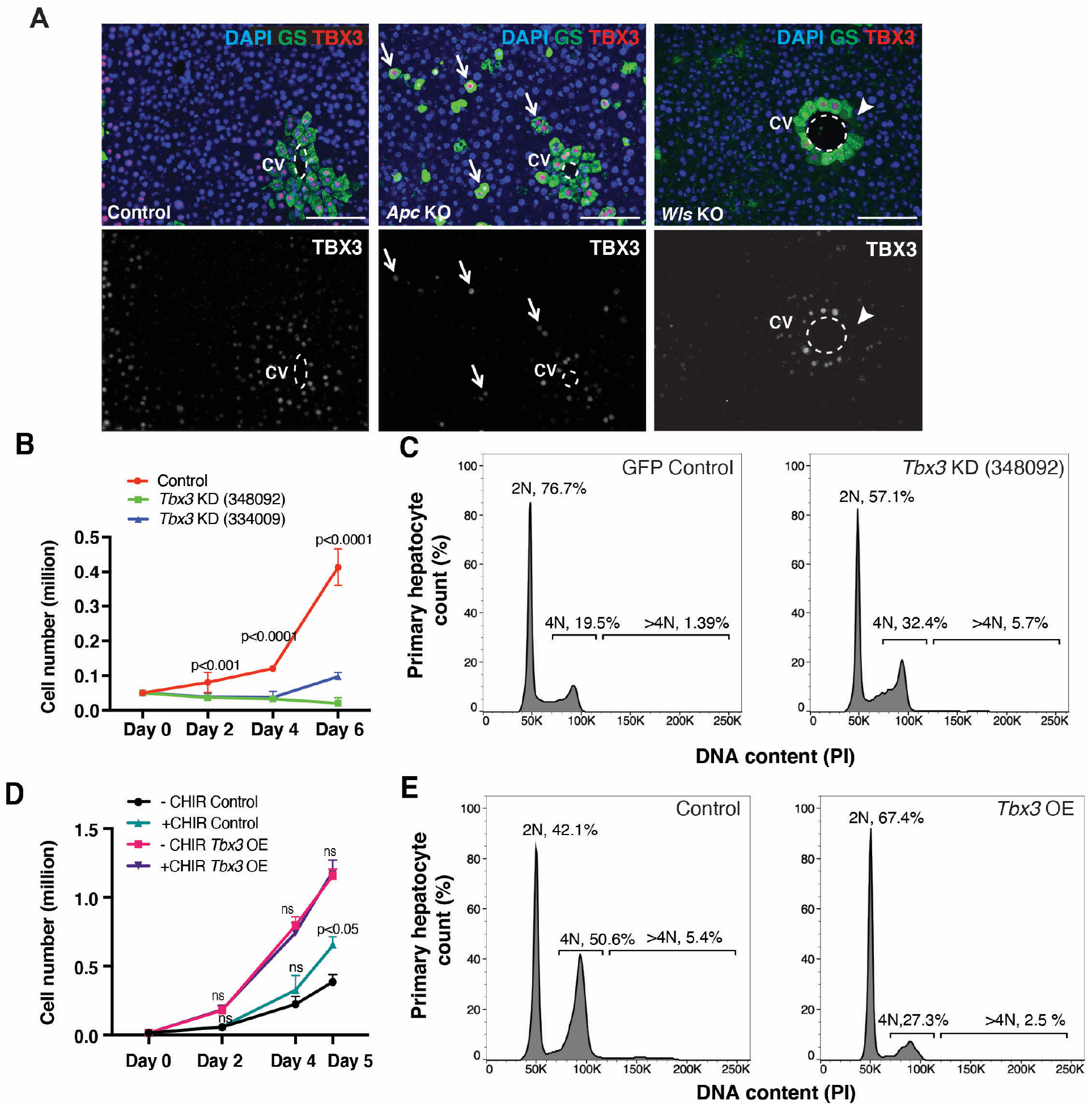
*Tbx3* acts downstream of Wnt signaling and promotes proliferation of cultured primary hepatocytes. (A) *Tbx3* expression is downstream of Wnt signaling. Immunofluorescence for GS and TBX3 is shown in control livers, CRISPR-Cas9 generated *Apc* KO clones, *Wls* KO; *(Ve-CadCreERT2; Wls*^*f/f*^) livers. Arrows: *APC* KO cells that express *Tbx3*. Arrowheads: *Wls* KO cells that do not express *Tbx3*. Scale bars, 100μm. Dashed lines delineate central veins. (B) *Tbx3* knock down (KD) slows down hepatocyte proliferation. Growth curves show the number of cells in Control (GFP) and two different *Tbx3* KD (shRNA #348092 and #334009) mouse primary hepatocyte cultures at days 0, 2, 4 and 6 after seeding. P-values represent comparisons between Control and each *Tbx3* KD condition from 3 independent experiments. (C) Tbx3 *KD* leads to increased numbers of polyploid hepatocytes in culture. Representative flow cytometry plots show ploidy distribution of *Control* and *Tbx3 KD* cells, stained with propidium iodide (PI). Percentages of ploidy classes are reported as averages from 3 independent experiments. (D) *Tbx3 OE* is sufficient to increase hepatocyte proliferation, in the absence of activated Wnt signaling. Growth curves of Control *(EF1α-GFP)* or *Tbx3* overexpressing (*Tbx3 OE*, EF1α-GFP-P2A-Tbx3) primary hepatocytes with or without added Wnt-activator CHIR99021(CHIR) are shown at days 0, 2, 4 and 5 after seeding. P-values represent comparisons between +CHIR control or *Tbx3* OE conditions and their respective -CHIR conditions. (E) *Tbx3* overexpression in primary hepatocytes leads to a higher percentage of diploid cells. Representative flow cytometry plots for ploidy distribution of Control and *Tbx3* OE hepatocytes, stained with PI. Percentages of ploidy distribution are averages from 3 independent experiments. Statistical significance was determined by Student’s t-test. Error bars represent standard deviations from 3 independent replicates. Scale bars, 100 μm. CV, central vein. KD, knock down. OE, overexpression. PI, propidium iodide. ns, not significant.

### Tbx3 controls ploidy of pericentral hepatocytes

We likewise found that TBX3 is required for ploidy control in hepatocytes within the growing postnatal liver. We generated inducible *Tbx3* loss-of-function mutant mice (*Tbx3* KO) carrying the *Axin2-rtTA; TetO-H2B-GFP*; *TetO-Cre*; *Tbx3* ^*f/f*^ construct. *Axin2rt-TA* is a doxycycline-inducible transgene which leaves the endogenous *Axin2* gene intact and is expressed in a pattern similar to *Tbx3* (Figure 2A) (Wang et al. 2015). To obtain maximal elimination of *Tbx3*, doxycycline was administered from postnatal week 2 to week 4 (Figure 2B). Nuclear size measurement (Figure 2C) and nuclear DNA content analysis (Figure 2D) revealed a significant decrease in the proportion of diploid nuclei in *Tbx3* KO livers (2N: 39.3% in Control vs 7.57% in Tbx3 KO) (Table S1) indicating that cells were able to complete S phase but not M phase. In parallel, we used the inducible and hepatocyte-specific *Albumin-CreERT2* (*Alb-CreERT2*) driver to eliminate *Tbx3* across all hepatocytes. We administered a single dose of tamoxifen at postnatal day 3 (P3) and analyzed the livers at different timepoints into adulthood (Figure 2E and Supplementary Figure 2, A and B). Loss of *Tbx3* did not affect liver shape or relative mass, at any timepoint (Figure S2, C and D). Consistent with the observations above, *Tbx3* KO livers were comprised of larger nuclei at all analyzed timepoints compared to control (Figure 2, F and G) and DNA content analysis confirmed increased ploidy in mutant tissues (Figure 2H and Table S1). Taken together, TBX3 is required to prevent polyploidization of hepatocytes *in vivo* and *in vitro*.

**Figure 2.**
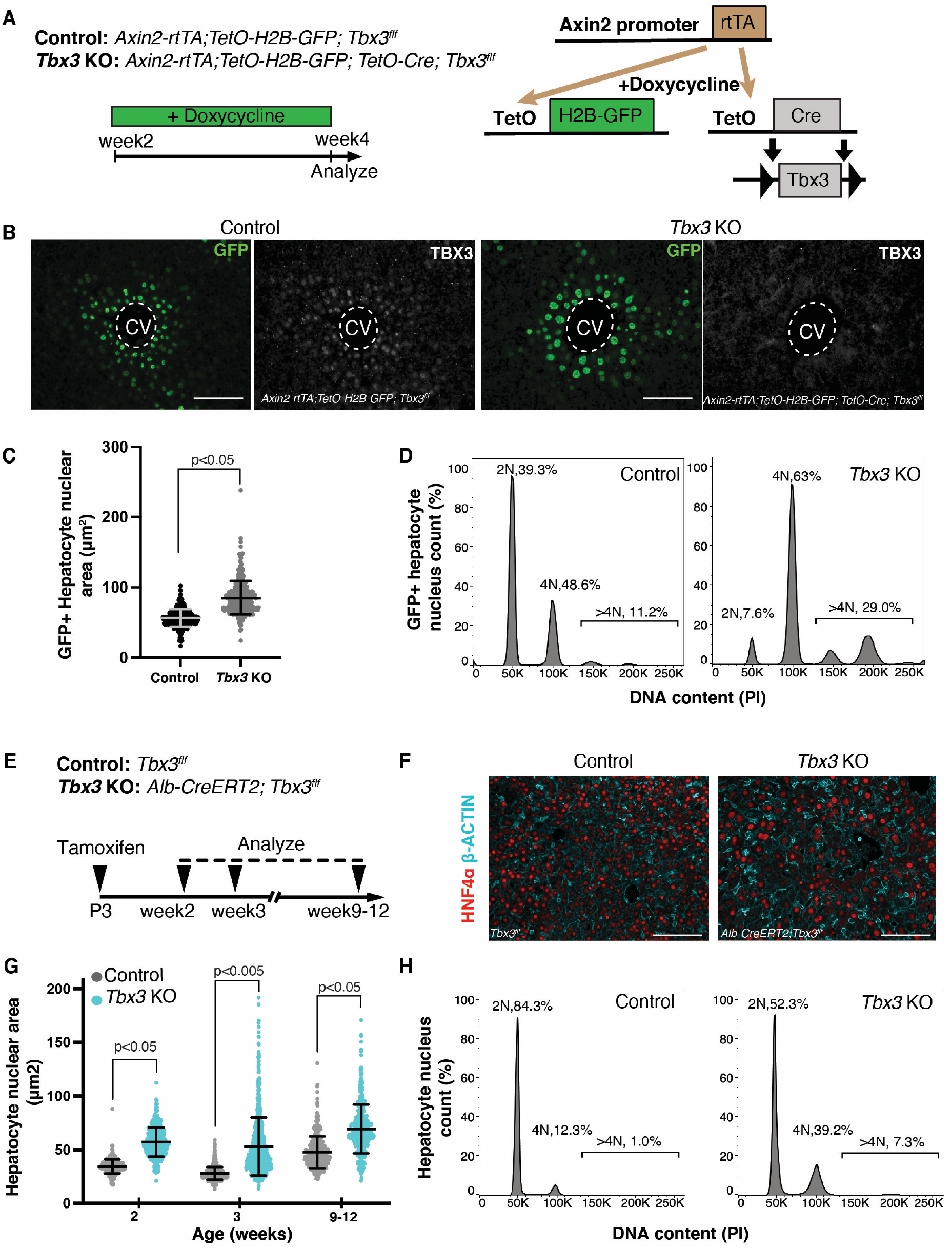
*Tbx3* ablation induces polyploidization of hepatocytes. (**A**) Schematic representation of mouse genotypes and experimental timeline. Animals were placed on a regimen of Doxycycline-treated drinking water from postnatal week 2 to week 4, to induce Axin2-rtTA-driven expression of GFP and deletion of *Tbx3* in pericentral hepatocytes (when the TetO-Cre transgene is present). (**B**) Axin2-rtTA-driven GFP expression and *Tbx3* deletion. Representative images of pericentral GFP+ nuclei detected by immunofluorescence, at postnatal week 4 in Control (*Axin2-rtTA; TetO-H2B-GFP; Tbx3*^*f/f*^) and *Tbx3* KO *(Axin2-rtTA; TetO-H2B-GFP; TetO-Cre; Tbx3*^*f/f*^) livers are shown. TBX3 immunofluorescence shows efficient deletion of the protein in *Tbx3* KO livers. (**C**) Axin2-rtTA-driven loss of *Tbx3* results in increased hepatocyte nuclear area. Distribution of GFP+ nuclear area at postnatal week 4 is shown. Bars indicate mean and standard deviation of all measured nuclei. T-test comparison with Welch’s correction was performed on the mean nuclear area of n=3 mice. (**D**) Axin2-rtTA-driven loss of *Tbx3* results in increased hepatocyte nuclear ploidy. Nuclear ploidy distribution of GFP+ hepatocytes in Control and *Tbx3* KO mice, stained with propidium iodide (PI) and measured by flow cytometry at postnatal week 4 is shown. Plots shown are chosen as representatives of each genotype from n=3 animals and noted percentages are averages. (**E**) Schematic representation of Alb-CreERT2-driven *Tbx3* deletion. Tamoxifen was administered to postnatal day 3 (P3) neonates with Control *(Tbx3*^*f/f*^) or *Tbx3* KO *(Alb-CreERT2; Tbx3*^*f/f*^) genotypes. Livers were harvested at postnatal weeks 2, 3 and 9-12 for analysis. (**F**) Visualization of hepatocyte membrane and nucleus in Alb-CreERT2-driven *Tbx3* KO livers. Representative HNF4α and β-ACTIN immunofluorescence images from Control and *Tbx3* KO livers at postnatal week 2 are shown. (**G**) Alb-CreERT2-driven loss of *Tbx3* results in increased hepatocyte nuclear area. Distribution of HNF4α+ nuclear area is shown. Bars indicate mean and standard deviation for all measured nuclei per genotype. T-test comparison with Welch’s correction was performed on the mean nuclear area from n=3 animals per genotype and timepoint. (**H**) Alb-CreERT2-driven loss of *Tbx3* results in increased hepatocyte nuclear ploidy. Nuclear ploidy distribution of hepatocytes stained with PI and measured by flow cytometry at postnatal week 2. Scale bars, 100 μm. CV, Central Vein. PV, Portal Vein. Dashed lines delineate central veins.

### TBX3 represses transcription of E2f7 and E2f8 to prevent hepatocyte polyploidization

To identify the possible mechanisms of TBX3 function, a known repressor of transcription, we investigated potential target genes using Chromatin immunoprecipitation-sequencing (ChIP-Seq) in *Tbx3*-overexpressing mouse primary hepatocytes. We found that TBX3 binds to promoter and enhancer regions of *E2f7* and *E2f8* and verified this interaction by ChIP-qPCR (Figure 3A and B). We hypothesized that Tbx3 represses *E2f7* and *E2f8*, and indeed *E2f7* and *E2f8* transcripts were both ectopically increased in *Tbx3* KO livers (Figure 3, C and D). Knock down and overexpression of *Tbx3* in cultured primary hepatocytes further corroborated the interactions with *E2f7/8* (Figure 3, E and F). To verify that repression of *E2f7* and *E2f8* by TBX3 occurs on the regulatory sequences of these genes, we employed a luciferase reporter assay in human hepatoblastoma HepG2 cells, which express *TBX3* at high levels (Feng et al. 2018). Addition of *E2f7* or *E2f8* enhancer regions to an HSV-TK reporter construct led to significant repression of the luciferase reporter activity compared to control vector with HSV-TK promoter only (Figure 3G). Moreover, reporter activity was increased when *TBX3* was knocked down, confirming that TBX3 is a specific transcriptional repressor of *E2F7* and *E2F8* (Figure 3G). In the adult human liver, *TBX3* is specifically expressed in pericentral hepatocytes, similar to the mouse liver (Figure S3A). Analysis of ChIP-Seq data from the ENCODE database showed that human TBX3 protein also binds to *E2F7* and *E2F8* enhancers (Figure S3B), suggesting that TBX3 may have conserved functions in both human and mouse livers.

**Figure 3.**
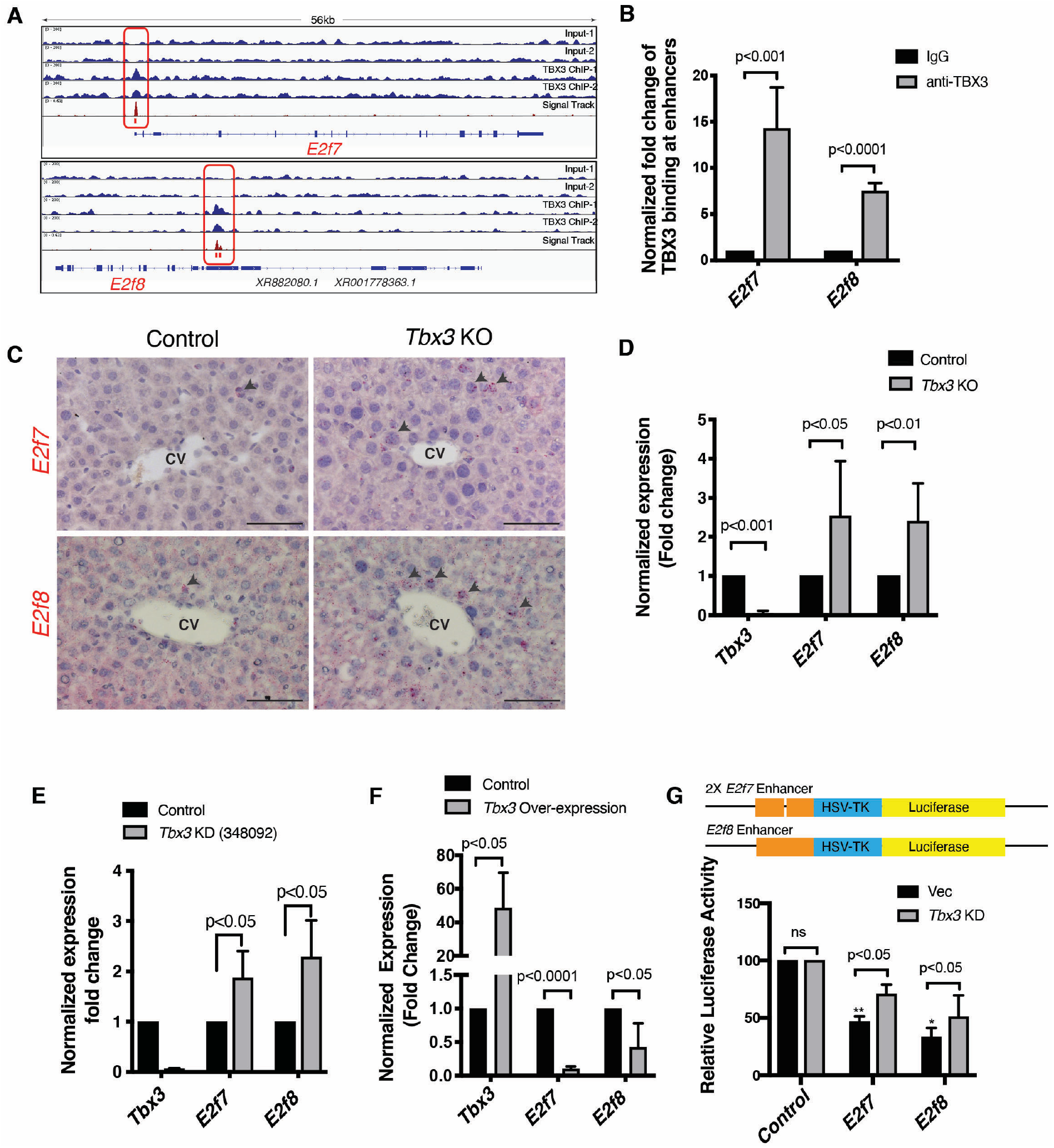
TBX3 directly represses transcription of *E2f7* and *E2f8*. (**A**) TBX3 binds to *E2f7* promoter and *E2f8 enhancer*, shown by Chromatin Immunoprecipitation-Sequencing (ChIP-seq) binding profiles (reads per million per base pair) for TBX3 at the *E2f7* and *E2f8 loci* in primary hepatocytes. Red boxes indicate significant binding peaks. These sites are tested in (B) and (G). Profiles are representative of two independent biological replicates. (**B**) Validation of binding sites outlined in (A) by ChIP-qPCR. **(C-E)** *E2f7* and *E2f8* are ectopically upregulated upon deletion of *Tbx3*. **(C)** *mRNA in situ* hybridization for *E2f7* and *E2f8* in *Control (Axin2-rtTA; TetO-H2B-GFP; Tbx3*^*f/f*^) and *Tbx3 KO (Axin2-rtTA; TetO-H2B-GFP; TetO-Cre; Tbx3*^*f/f*^) livers at postnatal week 4 show increased signal in the pericentral hepatocytes of *Tbx3 KO* livers. Arrowheads point to positive signals. **(D)** Quantitative real-time PCR (qRT-PCR) analysis of *Tbx3, E2f7* and *E2f8* transcripts from whole livers of the same samples as in (C) shows significantly increased levels of *E2f7* and *E2f8* in the absence of *Tbx3*. **(E)** qRT-PCR analysis shows increased *E2f7* and *E2f8* transcription in *Tbx3 KD* cultured mouse primary hepatocytes. (**F**) *E2f7* and *E2f8* are downregulated upon overexpression of *Tbx3*. qRT-PCR analysis of *Tbx3, E2f7* and *E2f8* transcripts in *Control (EF1α-GFP)* and *Tbx3 OE (EF1 α-GFP-P2A-Tbx3)* mouse primary hepatocytes. (**G**) TBX3 represses *E2f7* and *E2f8* expression by binding to their regulatory regions. Regions outlined in (A) as TBX3 binding sites were cloned into luciferase expression constructs for functional analyses in HepG2 cells and luciferase activity was measured. P-values noted via asterisks are comparisons of *Control* vs. *E2f7* and *Control* vs. *E2f8* in the Vector control condition. Scale bars, 100 μm. CV, Central Vein. KD, knockdown. Ns, not significant. Vec, Vector control. *P<0.05; **P < 0.01. Statistical significance was determined by Student’s t-test from n=3 independent experiments. Error bars indicate standard deviation.

The negative interactions between *Tbx3* and *E2f7*/*E2f8* and the up-regulation of *E2f7*/*E2f8* in the absence of *Tbx3* would imply that loss of all three genes would suppress the *Tbx3* knockout phenotype. To test this genetically, we generated triple conditional mutants of *Tbx3, E2f7* and *E2f8* (*Tbx3-E2f7-E2f8* TKO). First, we used the *Axin2-rtTA; TetO-GFP; TetO-Cre* system to delete all three genes by continuously administering Doxycycline from postnatal week 2 to week 4 (Figure 4 A). GFP-labeled nuclei in Tbx3-E2f7-E2f8 TKO livers were indistinguishable from controls in ploidy and size (Figure 4, B to D, and Table S1). Similar results were obtained in the pan-hepatocyte deletion model of Tbx3-E2f7-E2f8 TKO livers (Figure S4, A to C, and Table S1). These findings show that simultaneous removal of *E2f7, E2f8* and *Tbx3* resolves the polyploid phenotype caused by removal of *Tbx3* alone. These genetic interactions imply a linear pathway whereby Wnt activates *Tbx3*, which represses the mitotic inhibitors *E2f7* and *E2f8*; this therefore relieves cell cycle arrest and maintains hepatocytes in a diploid, dividing state.

**Figure 4.**
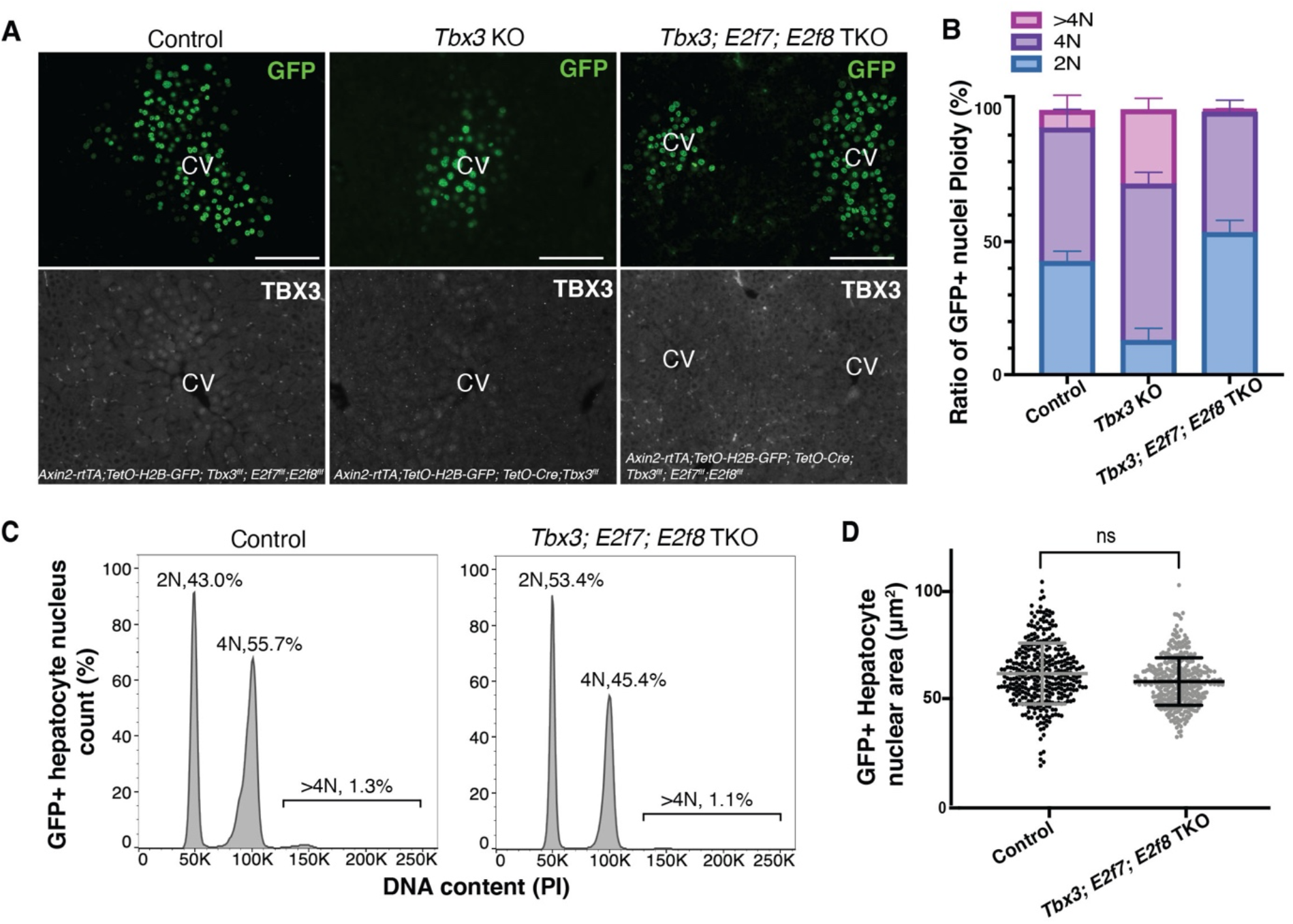
Genetic epistasis test showing TBX3 controls hepatocyte ploidy by repressing *E2f7* and *E2f8*. **(A)** Visualization of Axin2-rtTA-driven GFP expression in Control, *Tbx3* KO, *or Tbx3-E2f7-E2f8* TKO livers. GFP and TBX3 immunofluorescence in the pericentral zone in *Control (Axin2-rtTA; TetO-H2B-GFP; Tbx3*^*f/f*^; *E2f7*^*f/f*^; *E2f8*^*f/f*^*), Tbx3* KO *(Axin2-rtTA; TetO-H2B-GFP; TetO-Cre; Tbx3*^*f/f*^*) and Tbx3; E2f7; E2f8* TKO *(Axin2-rtTA; TetO-H2B-GFP; TetO-Cre; Tbx3*^*f/f*^; *E2f7*^*f/f*^; *E2f8*^*f/f*^) livers at postnatal week 4. **(B-D)** Deletion of *E2f7* and *E2f8* along with *Tbx3* restores the balance of ploidy. **(B)** Bar graph summarizes nuclear ploidy distribution of GFP+ nuclei from each genotype at postnatal week 4. **(C)** Nuclear ploidy plots of GFP+ hepatocytes in *Control* and *Tbx3-E2f7-E2f8* TKO mice, stained with Propidium Iodide (PI) and measured by flow cytometry. Plots shown are chosen as representatives of each genotype and noted percentages are averages from n=3 *Control* and n=4 *Tbx3-E2f7-E2f8* TKO mice. **(D)** Measurements of nuclear area from GFP+ pericentral hepatocytes in Control (n=3 mice) and *Tbx3-E2f7-E2f8* TKO (n=4 mice). Bars indicate mean and standard deviation. P=0.2249, t-test with Welch’s correction comparison of mean nuclear area from biological replicates. Scale bars, 100 μm. CV, Central Vein. ns, not significant.

## Discussion

In contrast to the extensive insight on the role of growth factors that initiate S phase of the cell cycle, little is known about extracellular signals that regulate M phase. High Wnt/βcatenin signaling activity during cell division has suggested a link between Wnt signaling and mitosis through unknown mechanisms (Benham-Pyle et al. 2016; Niehrs and Acebron 2012). In this work we have shown that in the liver, M phase is regulated by Wnt signals through the Wnt target transcription factor *Tbx3*, which in turn represses the mitotic inhibitors *E2f7* and *E2f8*. Interestingly, activating mutations in Wnt signaling components are common in liver cancer (Nusse and Clevers 2017; Wang et al. 2019) while *E2f7* and *E2f8* as well as polyploidy are known to suppress tumorigenesis (Kent et al. 2016; Moreno et al. 2021; Wilkinson et al. 2019; Zhang et al. 2018). These mitotic inhibitors are expressed in several other tissues besides the liver where they are also implicated in arresting the cell cycle (Matondo et al. 2018; Mizuno et al. 2019; Pandit et al. 2012; Qi et al. 2015). Whether Wnt signaling likewise regulates mitosis by suppressing *E2f7* and *E2f8* in other tissues and contexts is relevant to identifying the widespread functions of Wnts in development and disease (Nusse and Clevers 2017).

## Materials and Methods

### Animals

The Institutional Animal Care and Use Committee at Stanford University approved all animal methods and experiments. Wild type C57BL/6J mice, *Axin2-rtTA (B6*.*Cg-Tg(Axin2-rtTA2S*M2)7Cos/J) (Yu et al. 2007), TetO-H2B-GFP (Tg(tetO-HIST1H2BJ/GFP)47Efu/J)(Hadjantonakis and Papaioannou 2004)*, and *TetO-Cre (B6*.*Cg-Tg(tetO-cre)1Jaw/J), Wls* ^*f/f*^ *(129S-Wls*^*tm1*^.*1Lan/J)* (Carpenter et al. 2010) strains were obtained from The Jackson Laboratory (JAX). *Tbx3* ^*f/f*^ mice were a gift from Dr. Anne Moon (Frank et al. 2013). *Alb-CreERT2* mice were a gift from Dr. Julien Sage (Schuler et al. 2004). *E2f7* ^*f/+;*^ *E2f8* ^*f/+*^ were gifted by Dr. Alain de Bruin (Li et al. 2008) and were re-derived at the Stanford Transgenic Facility. *Cdh5-CreERT2* mice were used as previously described (Sorensen et al. 2009).

For knocking out *Tbx3* alone or *Tbx3*; *E2f7*; *E2f8* together using the *Axin2-rtTA* driver, animals were given 1 mg/ml doxycycline (Sigma D9891) in drinking water from postnatal day 14 (P14) until P28. In experiments involving the *Alb-CreERT2* driver, neonatal P3 mice received a single intragastric injection of 0.08mg tamoxifen (Sigma T5648), dissolved in corn oil with 10% ethanol.

For knocking out *Wls, Cdh5-CreERT2; Wls* ^*f/f*^ mice aged 8-10 weeks of aged received intraperitoneal injections of tamoxifen on four consecutive days and sacrificed at 7 days after last dose of tamoxifen.

All mice were housed with a 12-hour light/dark cycle with *ad libitum* access to water and normal chow.

### CRISPR/Cas9 mediated *Apc* deletion

Adult CRISPR/Cas9 knockin mice were obtained from JAX. A single intraperitoneal dosage of AAV-sgAPC was administered at a dose of 1×10^13 genome copies/kg and livers were collected for analysis 4 weeks after induction.

### Tissue Collection and processing

Livers were collected, fixed overnight at room temperature in 10% neutral buffered formalin, dehydrated, cleared in HistoClear (Natural Diagnostics), and embedded in paraffin. Sections were cut at 5μm thickness, de-paraffinized, re-hydrated and processed for further staining via immunofluorescence or *in situ* hybridization assays as described below.

### Immunofluorescence and Immunocytochemistry

Tissue Slides were subjected to antigen retrieval with Tris buffer PH=8.0 (Vector Labs H-3301) in a pressure cooker. They were then blocked in 5% normal donkey serum in PBS containing 0.1% Triton-X, in combination with the Avidin/Biotin Blocking reagent (Vector Labs SP-2001). Sections were incubated with primary and secondary antibodies and mounted in Prolong Gold with DAPI medium (Invitrogen). Biotinylated goat antibody (Jackson Immunoresearch 705-065-147) was applied to sections stained with Tbx3, before detection with Streptavidin-647. GS and b-Actin staining was performed with the Mouse-on-Mouse detection kit (Vector Labs) according to manufacturer’s protocol. The following antibodies were used: GFP (chicken, 1:500, Abcam ab13970), Tbx3 (goat, 1:50, Santa Cruz sc-17871), GS (mouse, 1:500; Millipore MAB302), b-Actin (mouse, 1:100; Abcam ab8226), Hnf4a (rabbit 1:50; Santa Cruz sc8987). Samples were imaged at 20X magnification using a Zeiss Imager Z.2 and processed and analyzed with the ImageJ software.

For immunocytochemistry, plated cells were fixed with 4% paraformaldehyde, blocked in 5% normal donkey serum in PBS containing 0.1% Triton-X and stained with primary and secondary antibodies as indicated above. Cells were imaged using a Zeiss Spinning Disk Confocal Microscope.

#### mRNA In Situ Hybridization

*In situs* were performed using the manual RNAscope 2.5 HD Assay-Red Kits (Advanced Cell Diagnostics) according to the manufacturer’s instructions. Images were taken at 20x magnification on a Zeiss Imager Z.2 and processed using the ImageJ software. Probes used in this study were *E2f7* (target region: 612-1526), *E2f8* (target region: 911-1893).

### RNA isolation and Real time quantitative PCR

Liver samples were homogenized in TRIzol (Invitrogen) with Pestle Motor Mixer (Argos Technologies A0001) or bead homogenizer (Sigma). Total RNA was purified using the RNeasy Mini Isolation Kit (Qiagen) and reverse-transcribed (High Capacity cDNA Reverse Transcription Kit, Life Technologies) according to manufacturer’s protocol. Quantitative RT-PCR were performed with TaqMan Gene Expression Assays (Applied Biosystems) on a StepOnePlus Real-Time PCR System (Applied Biosystems). Relative target gene expression levels were calculated using the delta-delta CT method as previously described(Livak and Schmittgen 2001). Gene Expression Assays used were *Gapdh* (Mm99999915_g1) as control, *Tbx3* (Mm01195719_m1), *E2f7* (Mm00618098_m1), and *E2f8* (Mm01204165_g1).

### Hepatocyte isolation and culture

Hepatocytes were isolated from 8-week-old mice by a two-step collagenase perfusion technique as previously described (Peng et al. 2018). 6-well plates (Greiner Bio-One 657160) were pre-coated with collagen (Corning 354236). Primary hepatocytes were plated into 2 ml of expansion media and media was replaced every 2-3 days. Basal media consisted of William E media containing 1% (v/v) Glutamax, 1% (v/v) Non-Essential Amino Acids, 1% (v/v) penicillin/streptomycin (all from Gibco), 0.2% (v/v) normocin (Invivogen), 2 % FBS (Omega), 2% (v/v) B27 (Gibco), 1% (v/v) N2 supplement (Gibco), 100 mM nicotinamide (Sigma-Aldrich), 1.25 mM N-acetylcysteine (Sigma-Aldrich), 10 μM Y27632 (Peprotech), 1 μM A83-01 (Tocris). Expansion media contained 3 μM CHIR99021 (Peprotech), 25 ng/mL EGF (Peprotech), 50 ng/mL HGF (Peprotech) and 100 ng/ml TNFα (Peprotech). To passage, cells were incubated with TrypLE Express (Gibco) for 5 minutes at 37 °C. Dissociated hepatocytes were transferred into new plates with fresh expansion media. Remaining cells were transferred to basal media and centrifuged at 300 g for 4 min. Cells were resuspended in Bambanker (Wako) and stored at −80 °C for be thawed following standard procedures for subsequent cultures.

### Lentiviral gene delivery to primary hepatocytes

At 24 hours prior to transfection, HEK293T cells were plated in 6-well plate at 8 × 105 cells per well in DMEM (10 % v/v FBS). Cells growing at ∼70-80 % confluency were transfected with 1 μg of pLKO.1-puro-CMV-TurboGFP (Sigma Aldrich) or one of five mouse *Tbx3* MISSION® shRNAs (TRCN0000348157, TRCN0000348092, TRCN0000095870, TRCN0000334009, TRCN0000095872) along with 2nd generation lentiviral packaging and envelope plasmids, 0.75 μg of psPAX2 and 0.25 μg of pMDL.g (gifts from D. Trono, Addgene #12260, #12259). At 36 hours post transfection, the media containing lentiviral particles was collected and passed through a 0.45 μm filter. Polybrene was added to a final concentration of 4 μg/ml. The filtered media containing lentivirus was added at a 1:1 (v/v) dilution to target primary hepatocytes. A second infection was performed at 60 hours post transfection. Hepatocytes were put on expansion media with puromycin (10 μl/ml) 1 day after the second transfection for 2 days for selection. Hepatocytes were harvested 4 days after the second transfection for qRT-PCR analysis and two out of five *Tbx3* shRNA were verified to be effectively knocking down *Tbx3*. 50,000 cells from these two lines were plated for growth curve analysis. Cell counts were performed using the Cellometer K2 (Nexelcom).

### Generation of Tbx3-HA ectopic expression vector

*Tbx3* was amplified from the MGC Fully Sequenced Mouse *Tbx3* cDNA (Ge Dharmacon MMM1013-202797681) with forward primer: 5’-TAAGCTTCTGCAGGTCGACTATGAGCCTCTCCATGAGAGATCC-3’and reverse primer: 5’-GAACATCGTATGGGTACATTGGGGACCCGCTGCAAGAC-3’and inserted into pKH3(addgene 12555) to add HA-tag at C-terminal of Tbx3 by NEBuilder® HiFi DNA Assembly Cloning Kit (E5520S). Then *Tbx3-HA* was amplified with forward primer: 5’ TGGAGGAGAACCCTGGACCTATGAGCCTCTCCATGAGAG-3’and reverse primer: TGATCAGCGGGTTTAAACTCAGCGTAATCTGGAACATC-3’and inserted into *pT2-SVNeo-EF1α-eGFP-P2A_PmeI* (modified from *pT2-SVNeo-EF1α-eGFP-P2A_EcoRI*, gift from Eric Rulifson).

### Applying Sleeping Beauty transposon system in primary hepatocytes

Primary hepatocytes were cultured with expansion media till 60% confluent. Sleeping Beauty system was applied as described in the plasmid DNA transfection protocol from TransIT-X2® Dynamic Delivery System (Mirus Biotech®) with modifications. 2.25 μg of *pT2-SVNeo-EF1α-eGFP or pT2-SVNeo-EF1α-eGFP-P2A-Tbx3-HA* along with the transposase in a ratio of 10:1 added with *Trans*IT-X2 and incubate 48 hours. Then cells were put on G418 selection with expansion media for 48 hours. Cells expressing GFP only or GFP together with Tbx3 were expanded for growth curve or flow cytometry analysis.

### Hepatocyte nuclei isolation and ploidy analysis

Hepatocyte nuclei preparation method was developed by modifying the chromatin preparation protocols described previously (Ghavi-Helm and Furlong 2012; Lemke et al. 2008). Liver lobes were homogenized in cold 1% formaldehyde in PBS with a loose pestle and dounce homogenizer with 15-20 strokes and fixed for 10 min at room temperature. Samples were incubated for 5 min with glycine at a final concentration 0.125 M and centrifuged at 300 g for 10 min, at 4°C. Pellets were washed in PBS and re-suspended with 10 ml cell lysis buffer (10 mM Tris-HCL, 10 mM NaCl, 0.5% IGEPAL) and filtered through 100 μm cell strainers. A second round of homogenization was performed by 15-20 strokes with a tight pestle. Nuclei were pelleted at 2000 g for 10 min at 4°C and re-suspended in 0.5ml PBS and 4.5 ml of pre-chilled 70% ethanol and stored at −20°C for ploidy analysis.

Nuclei were re-suspended in 5 ml of PBS and ca. 1 million nuclei were stained with 0.5 ml of FxCycle PI/RNase (Thermo Fisher, F10797) staining solution for 15-30 minutes at room temperature. Primary hepatocytes were fixed and stained the same way. Both nuclei and cells were analyzed on FACS ARIA II (BD). Data were processed with FACS Diva 8.0 software (BD) and FlowJo v10 (FlowJo). Doublets were excluded by FSC-W×FSC-H and SSC-W×SSC-H analysis. Single-stained channels were used for compensation and fluorophore minus one control was used for gating.

### Chromatin IP qPCR and Sequencing

Three to five million of *Tbx3* overexpressing primary hepatocytes were used for each chromatin IP reaction as recommended (Landt et al. 2012). Chromatin was prepared with truChIP™ Chromatin Shearing Reagent Kit (Covaris 520154) and sheared with Covaris S220 according to the kit manual. 4 μg anti-HA antibody (Roche Sigma 11867423001) was used for immunoprecipitation with BioVision immunoprecipitation kit (K286-25). Rat IgG1 isotype control (Thermo Fisher MA1-90035) was used for IgG control. Chromatin IP washing steps, Input and ChIP DNA preparation were modified as described previously (Ghavi-Helm and Furlong 2012) and sent for sequencing through NextSeq 500 system (Illumina). Sequencing data was mapped as previously described with modifications (Landt et al. 2012). Raw reads were mapped with Bowtie 2 (Langmead and Salzberg 2012) and processed and sorted with Samtools (Li et al. 2009). Peaks were called using MACS2 as previously described (Feng et al. 2011; Gaspar 2018; Zhang et al. 2008) with modifications from subcommands.

Purified Input, IgG and ChIP DNA following the chromatin immunoprecipitation were also used in ChIP-qPCR and calculated as described previously(Lemke et al. 2008). Forward:5’-AAGGTCATCCCAGAGCTGAA-3’ and Reverse: 5’-CTGCTTCACCACCTTCTTGA-3’ primers detect *Gapdh* promoter area were used as internal control. *E2f7* forward:5’-CGCCAGAGGAGGTGTTTTAG-3’ and reverse: 5’-CGGGCAGAGCGTGTAGAT-3’ or *E2f8* forward: 5’-GAACACTTGGGTGACCCTGA-3’ and reverse: 5’-CAAAGGGAATGCACACTGG-3’were used to detect *E2f7 or E2f8* promoter area that bound by Tbx3.

### Luciferase assay for promoter function

The HSV-TK promoter was cloned upstream of luciferase in the pGL4.10[luc2] plasmid (Promega). A 122 bp region of the *E2f7* promoter or a 2.1kb region of the *E2f8* enhancer was cloned into the XhoI site upstream of the HSV-TK promoter. HepG2 cells were co-transfected with 250 ng of the test plasmid and 250 ng of control pRL-TK plasmid, using the TransIT-X system (Mirus). Dual Luciferase Reporter assay (Promega) was performed at 48 hrs post-transfection with luminescence detected by a luminometer (Berthold Technologies). Firefly luciferase activity was normalized to Renilla luciferase in each sample.

### Quantification of nuclear area

Liver tissues were sectioned at 5μm, stained with anti-GFP or anti-HNF4α antibodies, and imaged at 20X magnification. Hepatocyte nuclei were detected automatically with the “Analyze particles” feature of Fiji (ImageJ) software to encircle GFP- or HNF4α-positive nuclei and selected regions were validated or corrected manually when necessary. Area of each encircled nucleus was then measured with the “Measure” feature. The number of nuclei measured per animal was as followed: *Control*=267-339, n=3mice, *Tbx3* KO=222-310, n=3 mice (Fig. 2C); *Control* week 2*=* 360-392, n=3mice, *Tbx3* KO week 2*=* 341-431, n=3mice, *Control* week 3*=* 436-714, n=3 mice, *Tbx3* KO week 3*=*298-714, n=3 mice, *Control* week 9-12*=278-* 411, n=3 mice, *Tbx3* KO week 9-12*=182-399*, n=3 mice (Fig. 2G), *Control=* 265-338, n=3 mice, *Tbx3-E2f7-E2f8* TKO*=* 303-370, n=4 mice (Fig. 4D), *Control=* 348-375, n=3 mice, *Tbx3* KO*=* 303-409, n=3 mice, and *Tbx3-E2f7-E2f8* TKO*=* 493-626, n=3 mice (Fig. S4C).

## Acknowledgments

We are grateful to WC. Peng for sharing primary hepatocytes culture protocol. We would like to thank C. Y. Logan, D. B. Azizoglu, and K. M. Loh for critically reading the manuscript and providing valuable insight, E.J. Rulifson for sharing reagents, the Stanford Functional Genomics Facility for ChIP-Seq library preparation and sequencing, and Wenyu Zhou at Bioinformatics-as-a-Service (BaaS) from Genetic Bioinformatics Service Center of Stanford School of Medicine for helping with ChIP-Seq analysis.

## Supplementary Figures

**Figure S1.**
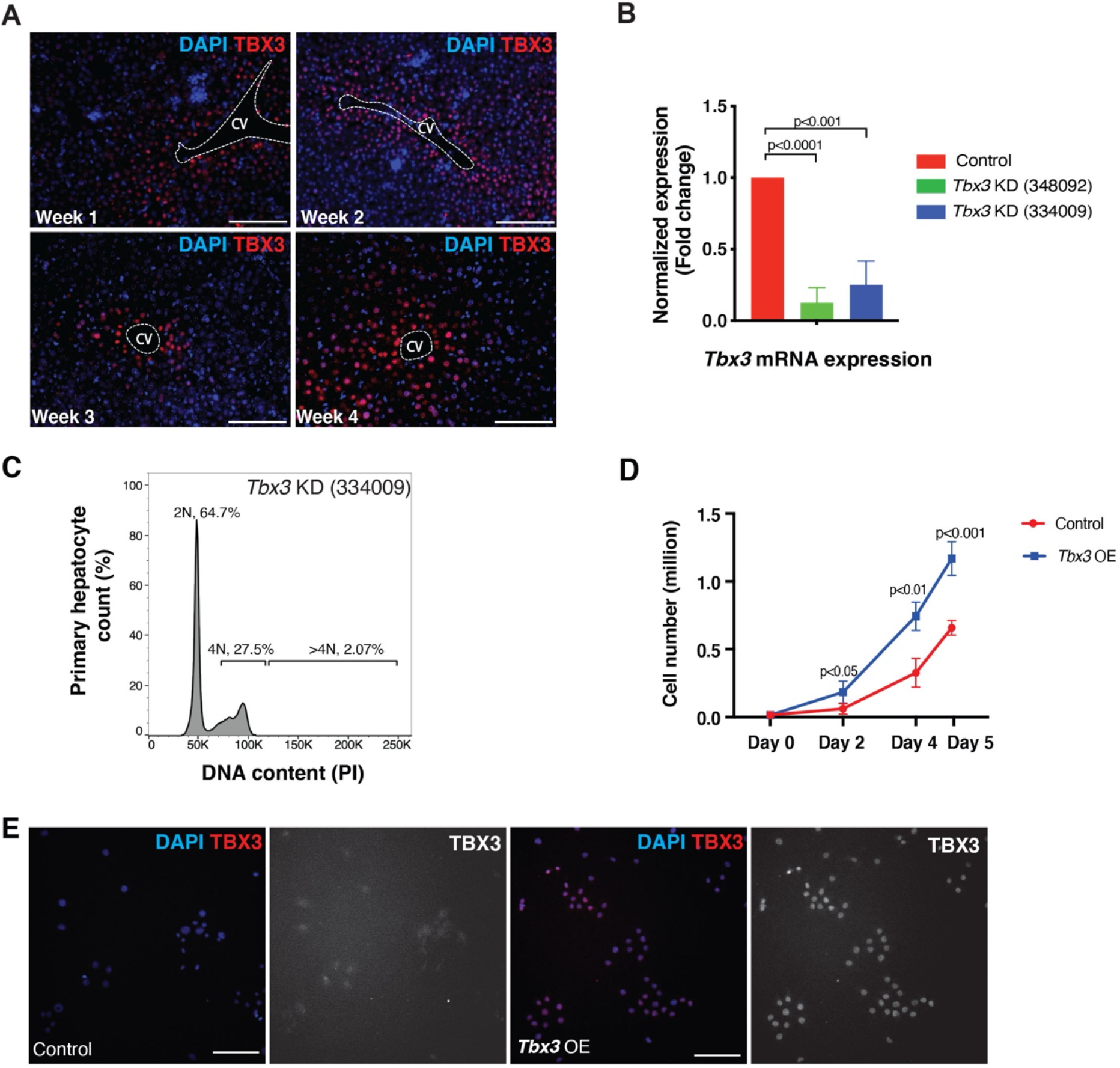
Expression of *Tbx3 in vivo* and its modulation *in vitro*. (**A**) *Tbx3* is expressed broadly in the pericentral zone during the postnatal growth phase. TBX3 protein is detected by immunofluorescence in hepatocytes surrounding the central veins, at postnatal weeks 1-4. Representative images are shown. (**B**) Efficient knockdown (KD) of *Tbx3* in hepatocytes. Cultured mouse primary hepatocytes were treated with control or one of two different *Tbx3* shRNA constructs (# #348092 and #334009). Normalized fold change in expression of *Tbx3* mRNA levels measured by qRT-PCR is shown. Statistical significance was determined by Student’s t-test between control and each *Tbx3* KD condition from n=3 independent experiments. (**C**) *Tbx3 KD* with shRNA #334009 resulted in a minor increase in the percentage of polyploid cells, compared to Control in Fig. 1C. Representative flow cytometry plots show ploidy distribution of *Tbx3* KD cells, stained with propidium iodide (PI). Percentages of ploidy classes are reported as averages from n=3 independent experiments. (**D**) *Tbx3* Overexpression (OE) leads to increased hepatocyte proliferation in culture. Growth curves of Control *(EF1α-GFP)* and *Tbx3* overexpressing (*Tbx3 OE*, EF1α-GFP-P2A-Tbx3) primary hepatocytes are shown at days 0, 2, 4 and 5 after seeding. Statistical significance was determined by Student’s t-test from n=3 independent experiments. (**E**) Cultured *Tbx3* overexpressing primary hepatocytes. Representative TBX3 immunofluorescence images of Control or *Tbx3* OE mouse primary hepatocytes are shown. Scale bars, 100μm. CV, central vein. KD, knock down. OE, overexpression. PI, propidium iodide. Dashed lines delineate central veins.

**Figure S2.**
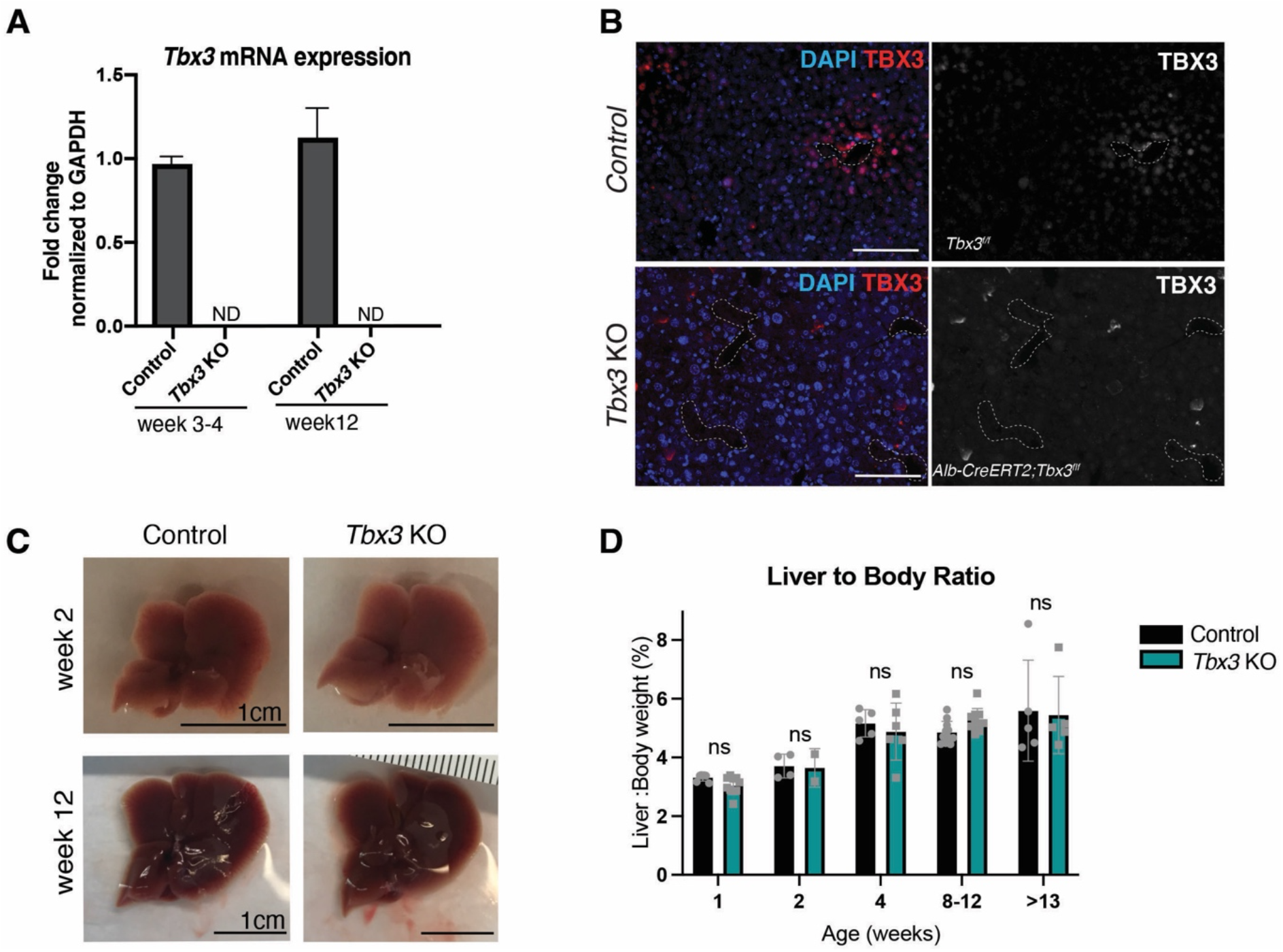
AlbCreERT2-driven loss of *Tbx3* does not impact liver size or morphology. (**A**) Efficient deletion of *Tbx3* mRNA with the Alb-CreERT2 driver. Neonates received a single dose tamoxifen at postnatal day 3 and Tbx3 deletion was verified by qRT-PCR in Control *(Tbx3*^*f/f*^) and *Tbx3* KO *(Alb-CreERT2; Tbx3*^*f/f*^) livers at postnatal weeks 3-4 and 12. (**B**) Efficient deletion of TBX3 protein with the Alb-CreERT2 driver. Representative images of TBX3 immunofluorescence, at postnatal week 3 in Control *(Tbx3*^*f/f*^) and *Tbx3* KO *(Alb-CreERT2; Tbx3*^*f/f*^) livers are shown. (**C**) Photographs of Control *(Tbx3*^*f/f*^) and *Tbx3* KO *(Alb-CreERT2; Tbx3*^*f/f*^) livers at postnatal weeks 2 and 12 show no changes to liver morphology. (**D**) Liver to body weight ratios at various weeks after birth show no significant differences in liver size between Control *(Tbx3*^*f/f*^) and *Tbx3* KO *(Alb-CreERT2; Tbx3*^*f/f*^) animals. Each data point is representative of one animal. ND, Not Detected. ns= not significant. Dashed lines delineate veins.

**Figure S3.**
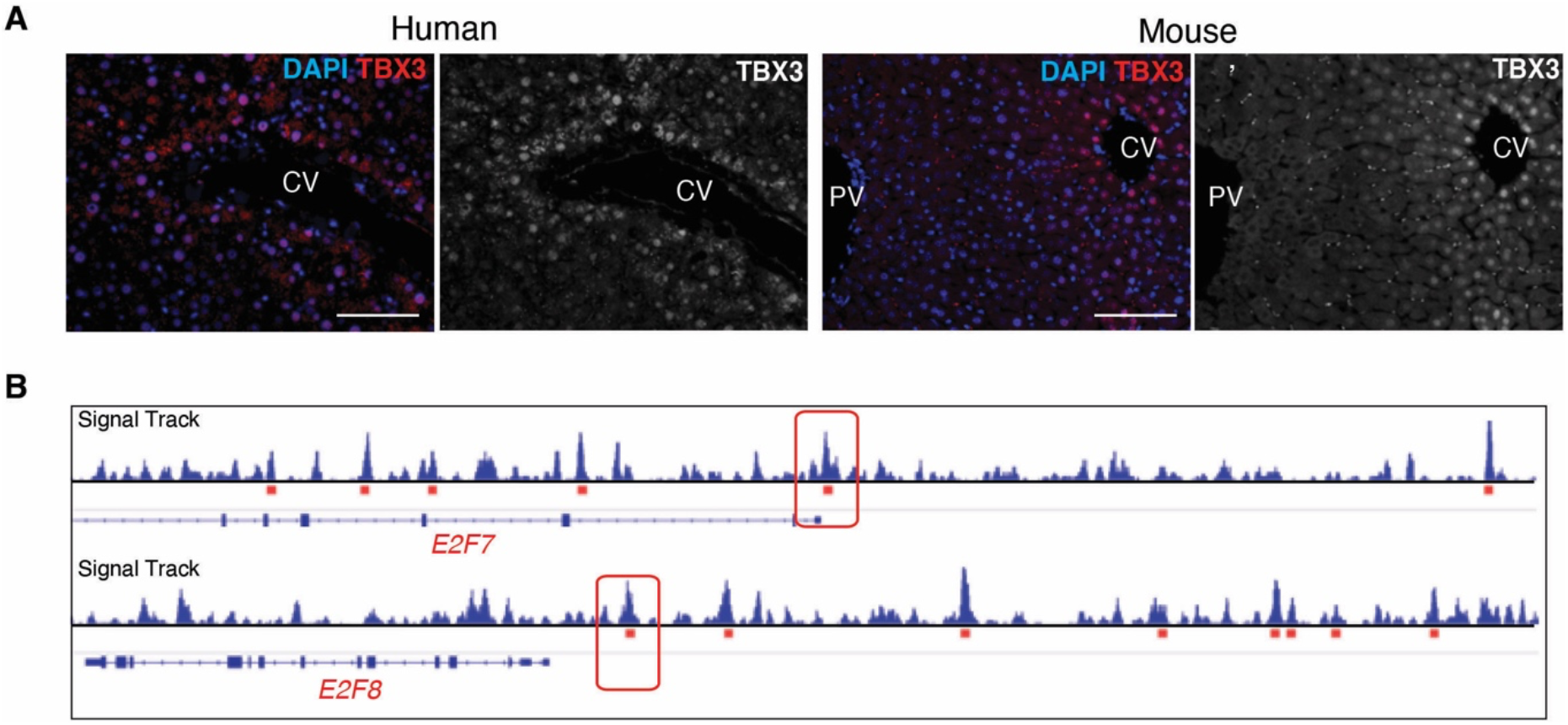
*Tbx3* is expressed in pericentral hepatocytes of the human liver and binds to *E2f7/E2f8* in human hepatocytes. (**A**) *Tbx3* is expressed in the pericentral hepatocytes of human livers. TBX3 protein is detected by immunofluorescence in both human and mouse livers. (**B**) TBX3 binds to *E2F7* and *E2F8* in human hepatocytes. TBX3 Chromatin Immunoprecipitation-Sequencing (ChIP-Seq) Signal Track from HepG2 Cells (ENCODE) for *E2F7* and *E2F8* loci. Regions underlined in red indicate significant binding peaks. Boxed regions are significant binding regions conserved in mouse. CV, Central Vein. PV, Portal Vein. Scale bars, 100 μm.

**Figure S4.**
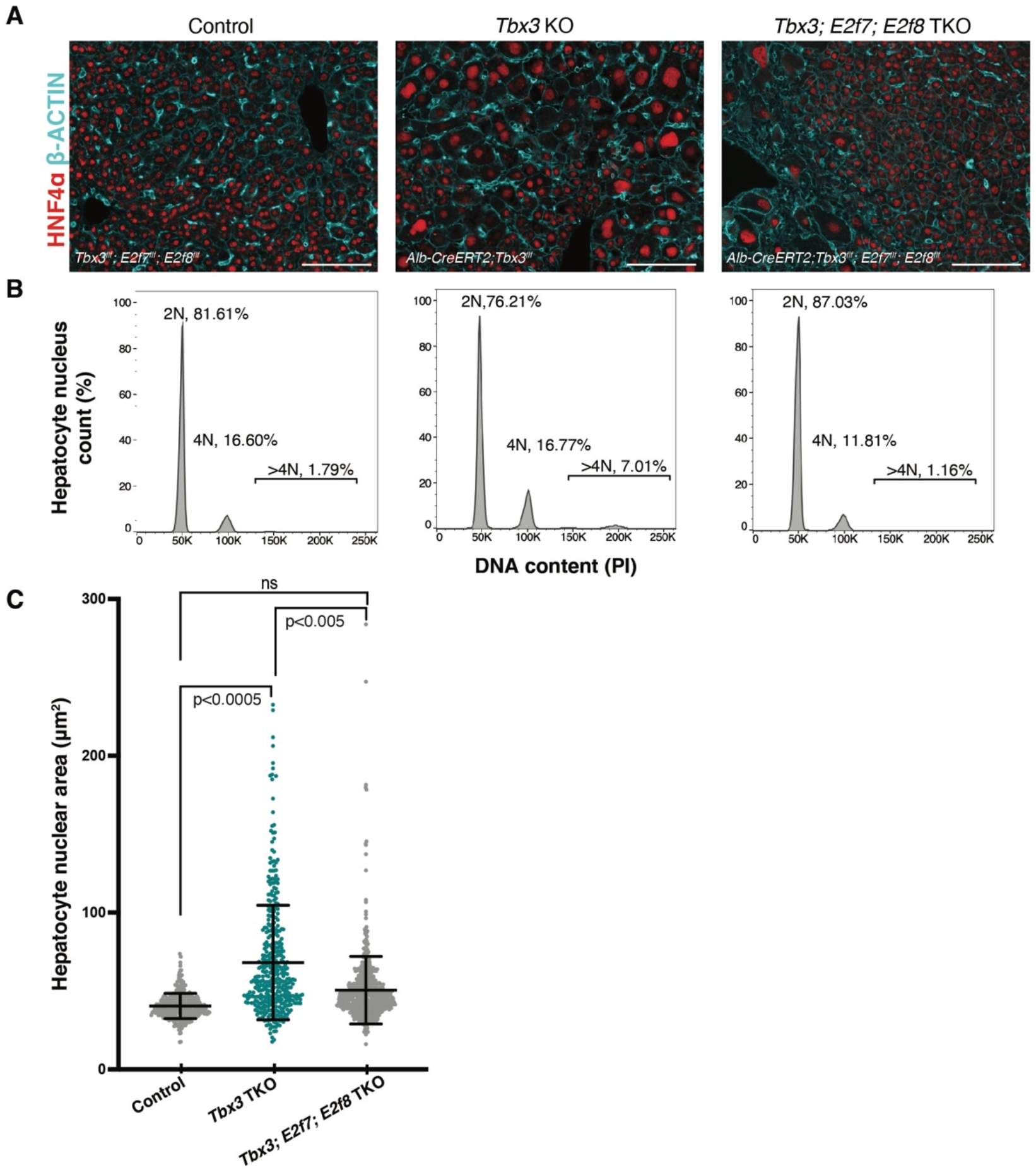
Genetic epistasis test showing TBX3 controls hepatocyte polyploidy by repressing *E2f7* and *E2f8*. **(A)** Membrane and nuclear visualization of Control or Alb-CreERT2-driven *Tbx3* KO or *Tbx3-E2f7-E2f8* TKO livers. Representative HNF4α and β-ACTIN immunofluorescence images at postnatal week 4 are shown. **(B-C)** Deletion of *E2f7* and *E2f8* along with *Tbx3* rescues the polyploidy phenotype of *Tbx3* KO. **(B)** Representative flow cytometry plots of nuclear ploidy distribution from n=5 Control *(Tbx3*^*f/f*^; *E2f7*^*f/f*^; *E2f8*^*f/f*^), n=4 *Tbx3* KO *(Alb-CreERT2; Tbx3*^*f/f*^) and n=4 *Tbx3-E2f7-E2f8* TKO *(Alb-CreERT2; Tbx3*^*f/f*^; *E2f7*^*f/f*^; *E2f8*^*f/f*^) livers at postnatal week 4. **(C)** Measurements of hepatocyte nuclear area in livers from (A). n=3 mice pre genotype. Bars indicate mean and standard deviation for all measured nuclei per genotype. Means of nuclear area were calculated for each animal. Means for each genotype were compared by one-way ANOVA. Ns= not significant (adjusted P=0.0626). Scale bars, 100μm. PI, propidium iodide.

**Table S1.** Average ploidy distribution of hepatocytes nuclei.

| Genotype | Average |  |  | p-Value |  |  |
| --- | --- | --- | --- | --- | --- | --- |
|  | 2N% | 4N% | >4N% | 2N% | 4N% | >4N% |
| *Control (GFP)* | 76.7 | 19.5 | 1.4 | p < 0.001 | p < 0.05 | p < 0.05 |
| *Tbx3 KD (shRNA #348092)* | 57.1 | 32.4 | 5.7 |  |  |  |
| *Control (EF1α-GFP)* | 42.1 | 50.6 | 5.4 | p < 0.001 | p < 0.001 | p < 0.05 |
| *Tbx3 overexpressing (Tbx3 OE, EF1α-GFP-P2A-Tbx3)* | 67.4 | 27.3 | 2.5 |  |  |  |
| *Axin2-rtTA; TetO-H2B-GFP; Tbx3<sup>fl/fl</sup>* | 39.3 | 48.6 | 11.2 | p < 0.001 | p < 0.05 | p < 0.005 |
| *Axin2-rtTA; TetO-H2B-GFP; TetO-Cre; Tbx3<sup>fl/fl</sup>* | 7.6 | 63 | 29 |  |  |  |
| *Axin2-rtTA; TetO-H2B-GFP; Tbx3<sup>fl/fl</sup>; E2f7<sup>fl/fl</sup>; E2f8<sup>fl/fl</sup>* | 43.0 | 55.7 | 1.3 | ns | ns | ns |
| *Axin2-rtTA; TetO-H2B-GFP; TetO-Cre; Tbx3<sup>fl/fl</sup>; E2f7<sup>fl/fl</sup>; E2f8<sup>fl/fl</sup>* | 53.4 | 45.4 | 1.1 |  |  |  |
| *Tbx3<sup>fl/fl</sup>* | 84.3 | 12.3 | 1.0 | p < 0.05 | p < 0.05 | ns |
| *Alb -CreERT2; Tbx3<sup>fl/fl</sup>* | 52.3 | 39.2 | 7.3 |  |  |  |
| *Tbx3<sup>fl/fl</sup>; E2f7<sup>fl/fl</sup>; E2f8<sup>fl/fl</sup>* | 81.6 | 16.6 | 1.8 | ns | ns | ns |
| *Alb -CreERT2; Tbx3<sup>fl/fl</sup>; E2f7<sup>fl/fl</sup>; E2f8<sup>fl/fl</sup>* | 87.0 | 11.9 | 1.2 |  |  |  |

